# Pervasive loss of regulated necrotic cell death genes in elephants, hyraxes, and sea cows (*Paenungualta*)

**DOI:** 10.1101/2024.04.04.588129

**Authors:** Meaghan Birkemeier, Arianna Swindle, Jacob Bowman, Vincent J. Lynch

## Abstract

Gene loss can promote phenotypic differences between species, for example, if a gene constrains phenotypic variation in a trait, its loss allows for the evolution of a greater range of variation or even new phenotypes. Here, we explore the contribution of gene loss to the evolution of large bodies and augmented cancer resistance in elephants. We used genomes from 17 Afrotherian and Xenarthran species to identify lost genes, i.e., genes that have pseudogenized or been completely lost, and Dollo parsimony to reconstruct the evolutionary history of gene loss across species. We unexpectedly discovered a burst of gene losses in the Afrotherian stem lineage and found that the loss of genes with functions in regulated necrotic cell death modes was pervasive in elephants, hyraxes, and sea cows (*Paenungulata*). Among the lost genes are *MLKL* and *RIPK3*, which mediate necroptosis, and sensors that activate inflammasomes to induce pyroptosis, including *AIM2*, *MEFV*, *NLRC4*, *NLRP1*, and *NLRP6*. These data suggest that the mechanisms that regulate necrosis and pyroptosis are either extremely derived or potentially lost in these lineages, which may contribute to the repeated evolution of large bodies and cancer resistance in Paenungulates as well as susceptibility to pathogen infection.

## Introduction

Gene duplication has long been known to have a prominent role in generating the molecular diversity that underlies phenotypic differences between species; for example, by generating functional redundancy that is beneficial because increased copy number (dosage) has beneficial effects, or redundancy that can resolve through the origins of proteins with new functions (neofunctionalization). Gene loss, in contrast, is most often thought to result from environmental or behavioral changes that lead to the loss of selection to maintain gene functions, such as the loss of bitter and sweet taste receptors in blood- and insect-eating bats (Hong and Zhao, 2014; Jiao et al., 2021; LU et al., 2021; Zhao et al., 2010). However, gene loss itself can directly lead to phenotypic differences between species when a gene constrains the range of evolutionarily relevant phenotypic variation in a trait; in these cases, gene loss breaks ancestral constraints on phenotypes, allowing traits to evolve beyond their ancestral ranges or for entirely new traits to evolve. For example, gene loss in yeast has been shown to increase evolvability by facilitating the emergence of a more diverse array of phenotypes, including multicellularity (Montrose et al., 2024a), while the loss of the heat shock protein Hsp90 chaperon reveals hidden phenotypic variation upon which selection can act in both *Drosophila* (Rutherford and Lindquist, 1998) and *Arabidopsis* (Queitsch et al., 2002). Similarly, the loss of *AMPD3* in sperm whales may have allowed for the evolution of long-diving times, while the loss of *SLC22A12* (URAT1), *SLC2A9* (GLUT9), and *SLC22A6* (OAT1) may have facilitated the evolution of fruit-based diet in bats (Sharma et al., 2018). Thus, gene loss can reveal previous hidden morphological variation and break ancestral developmental constraints, leading to the evolution of novel phenotypes (Montrose et al., 2024b).

Among the constraints on evolving exceptionally large bodies and long lifespans is an increased risk of developing cancer. If cells in large and small organisms have a similar risk of cancerous transformation and equivalent cancer suppression mechanisms, for example, organisms with many cells should have a higher prevalence of cancer than organisms with fewer cells (Caulin and Maley, 2011; Nagy et al., 2007; Peto, 2015a). Consistent with these expectations, there is a positive correlation between cancer prevalence, body size, and lifespan *within* species. For example, cancer prevalence increases with increasing adult height in humans (Benyi et al., 2019; Green et al., 2011; Nunney, 2018; Zhou et al., 2022) and with increasing body size in dogs (Dobson, 2013). However, there is no positive correlation between body size and cancer risk between mammalian species (Abegglen et al., 2015; Bulls et al., 2022; Compton et al., 2023); this lack of correlation is called ‘Peto’s Paradox’ (Peto, 2015b, 1975). Furthermore, there is a statistically significant *negative* correlation between body mass and cancer prevalence in Eutherian mammals (Bulls et al., 2022), indicating that mammals with enormous bodies have evolved particularly effective anticancer mechanisms.

The resolution to Peto’s paradox is simple – species with exceptionally large bodies have evolved enhanced cancer resistance. Numerous mechanisms have been proposed to explain the evolution of reduced cancer risk, including the gain of additional tumor suppressor genes and the loss of oncogenes. We have previously shown, for example, that duplication of genes with tumor suppressor functions was common in Afrotherians and Xenarthrans, especially in cancer-resistant species such as elephants and armadillos (Vazquez et al., 2022; Vazquez and Lynch, 2021). Here, explore whether gene loss in *Afrotheria* and *Xenarthra* is also associated with cancer biology using gene loss data inferred with the Tool to infer Orthologs from Genome Alignments (TOGA) from 17 species (Kirilenko et al., 2023). Surprisingly, we found genes that are essential for several regulated cell death modes have been lost in many Afrotherian lineages, including *MLKL* and *RIPK3*, which mediate necroptosis and have been lost in Paenungulates, and multiple sensor proteins that form inflammasomes to induce pyroptosis including *AIM2* in *Afrotheria*, *MEFV* in elephants, *NLRC4* in Paenungulates and *NLRP6* in hyraxes. These data suggest that several Afrotherian lineages have lost genes that directly mediate regulated necrotic cell death modes, which may contribute to the evolution of cancer resistance at the cost of susceptibility to microbial and viral infections.

## Results

### Identification of gene losses in *Atlantogenata*

We used gene loss data for 17 species (**Figure 1**), which was inferred with TOGA (Kirilenko et al., 2023) from 261 mammals in the Zoonomia alignments; lost genes are those that have been deleted from the genome or that have acquired inactivating mutations. We coded gene presence/absence data as binary discrete character states such that genes present in the genome of each species were coded as state 1, whereas genes lost were coded as state 0. As a preliminary analysis, we explored the structure of gene loss data with two complementary methods: 1) Fuzzy C-Means (FCM) clustering, a “soft” clustering approach that allows samples to have proportional membership in multiple clusters; and 2) Multidimensional scaling (MDS), a hard clustering approach in which samples can only belong to one group that is inferred by k-means clustering. Both approaches grouped species by taxonomic relationships, with greatest FCM cluster membership grouping *Paenungulata*, *Afroinsectiphilia*, and *Xenarthra* (**Figure 2A**), while MDS clustered species by superorder, i.e., *Afrotheria* and *Xenarthra*, along dimension 1 and *Afrotherian* grandorder, i.e., *Afroinsectiphilia* and *Paenungulata*, or *Xenarthran* order, i.e., *Cingulara* and *Pilosa*, along dimension 2 (**Figure 2B**). These data indicate that gene loss events cluster species by phylogenetic relatedness rather than randomly, as would be expected if there was no phylogenetic signal in gene loss data and they resulted from technical errors or sequencing artifacts. The Southern three-banded armadillo, however, was inferred to be its own cluster by both FCM and MDS, suggesting that this species has an unusual pattern of gene losses that may be artifactual. Thus, we conclude gene loss data may have phylogenetic signals resulting from patterns of shared derived (synapomorphic) loss events.

**Figure 1.**
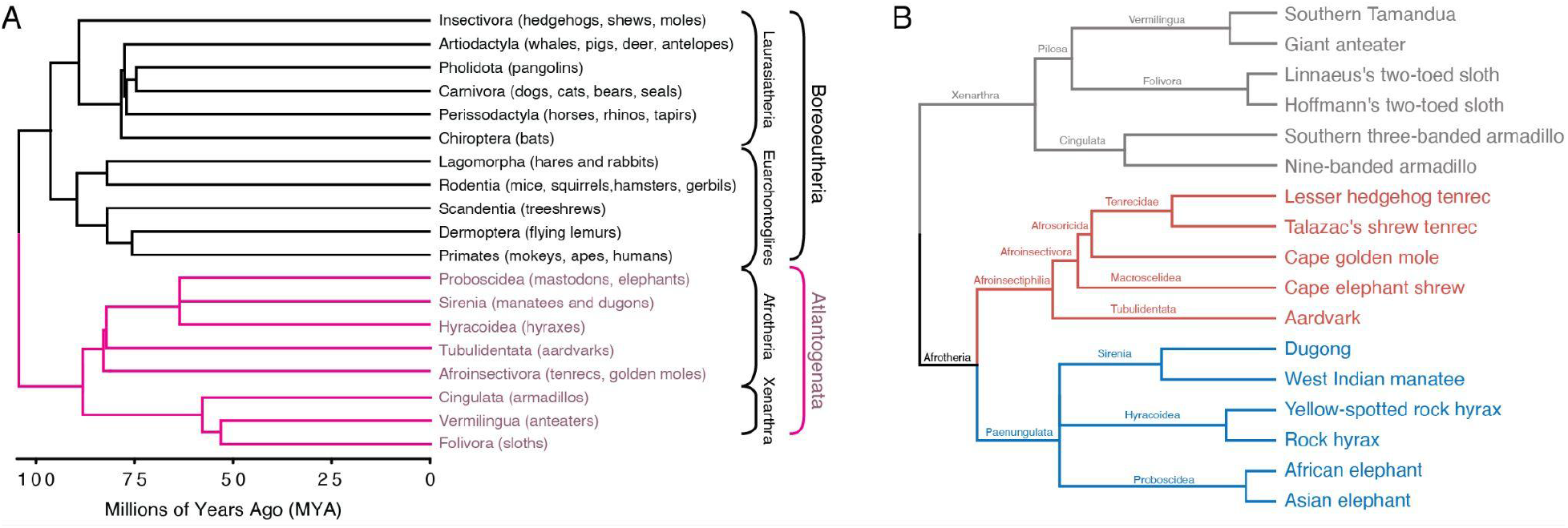
Eutherian mammal phylogenetic relationships. A. Phylogenetic relationships between Eutherian orders; examples of each order are given in parenthesis. Horizontal branch lengths are proportional to the time since divergence between lineages. Major clade names are shown; Atlantogenta are shown in pink. B. Phylogenetic relationships between Atlantogenatans. Note that horizontal branch lengths are arbitrary; the order *Xenarthra* is colored grey, *Paenungulata* is colored blue, and Afroinsectiphilia is colored red.

**Figure 2.**
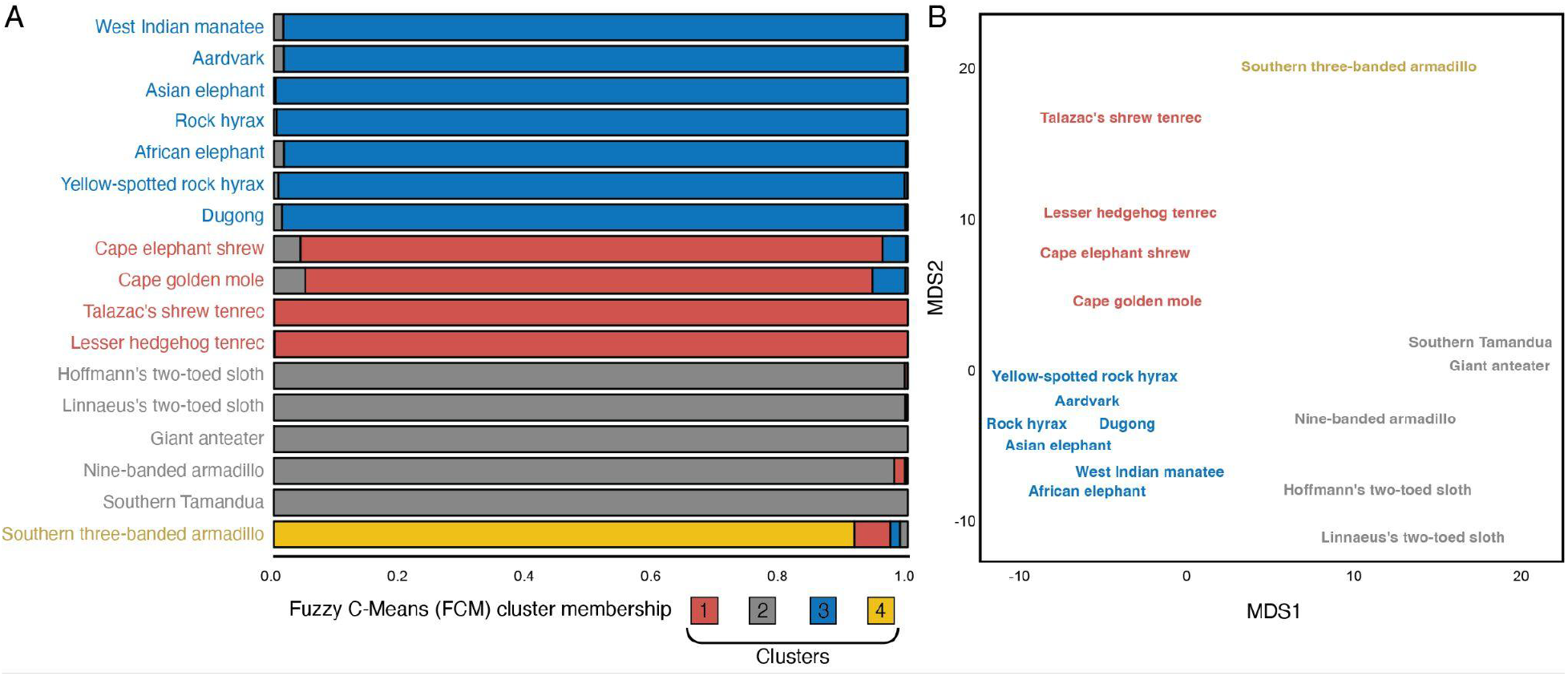
Gene loss data cluster species by phylogenetic relationships. A. Fuzzy C-Means cluster membership proportions of each species based on gene loss data. The degree of cluster membership is shown as a 100% stacked bar and colored according to the proportion of membership in each cluster (K=4). B. Multidimensional scaling (MDS) plot of species based on gene loss data. Cluster membership was inferred with K-means clustering (K=4).

### Gene loss data has a significant phylogenetic signal

We used the Dolpenny program in Phylip to infer the most parsimonious phylogenetic tree for the gene loss data under the Dollo criteria and using a "branch and bound" tree search algorithm; unlike regular parsimony, which minimizes the number of evolutionary events, the Dollo model allows only one forward change (0→1), but as many reversions (1→0) as necessary to explain the distribution of character states among extant species. Thus, the criteria of Dollo parsimony is to minimize the number of 1→0 reversions, which is the ideal model for reconstructing the evolution of gene loss events across a phylogeny because, once lost, a homologous gene can never reevolve (Sharma et al., 2018). Branch support was assessed with 100 bootstrap replicates and a parsimony variant of the Shimodaira-Hasegawa (SH) test that compared the parsimony score of the inferred most parsimonious tree to one in which each branch was individually collapsed to a polytomy. The most parsimonious phylogeny had low bootstrap support at deeper nodes but high SH-test support (*P*<0.000) for all branch bipartitions (**Figure 3A**) and generally recapitulated the species phylogeny with only three discordant relationships (**Figure 3B**): 1) Armadillos were not monophyletic and nested within *Folivora*; 2) Aardvark is placed as a sibling species to *Paenungulata* rather than as an outgroup to the *Afroinsectivora*; and 3) Tenrecs were not monophyletic. Correcting the phylogenetic placement of discordant taxa led to significantly worse parsimony scores, with an SH-test *P*<0.0001. These data indicate that there is phylogenetic signal in the pattern of gene losses in *Atlantogenata* and likely also convergent gene losses or systematic error in gene loss inference driving the discordant phylogenetic relationships between some species. Thus, we conclude that (the majority) of gene losses are likely real rather than false positives, i.e., genes erroneously inferred to be lost because of systematic biases in the TOGA method or genome assembly errors.

**Figure 3.**
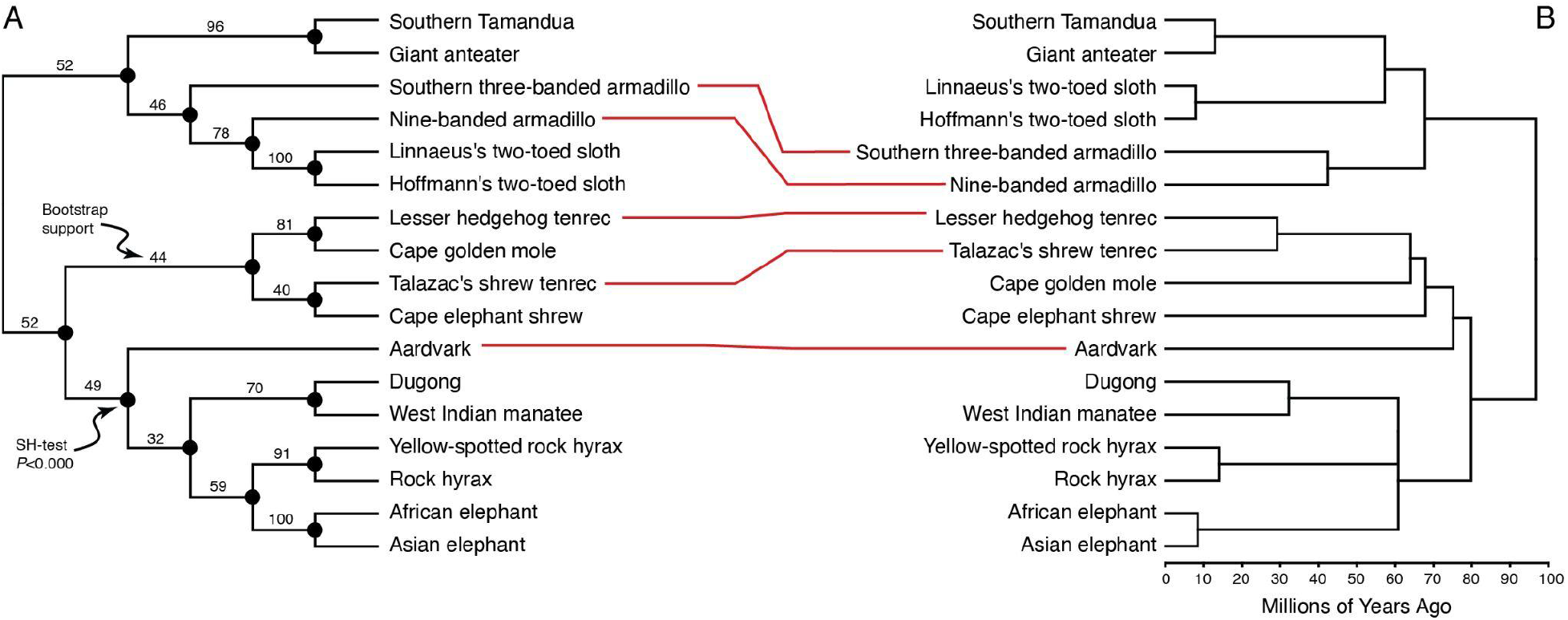
Parsimony tree inferred from gene loss data under the Dollo model reflects phylogenetic relationships. A. The most parsimonious phylogenetic tree for the gene loss data was inferred under the Dollo model using a "branch and bound" search. Branch support was assessed with 100 bootstrap replicates and a parsimony variant of the Shimodaira-Hasegawa (SH) test that compared the parsimony score of the inferred most parsimonious tree to one in which each branch was individually collapsed to a polytomy. Bootstrap support values are shown for each internal branch; all nodes ahd high SH-test support (*P*<0.000). B. The Parsimony tree reflects the species phylogeny with only three discordant relationships (red lines): 1) Armadillos were not monophyletic and nested within *Folivora*;. 2) Aardvark is placed as a sibling species to *Paenungulata* rather than as an outgroup to the *Afroinsectivora*; and 3) Tenrecs were not monophyletic. **Figure 3 – source data 1. Nexus file with gene loss matrix and phylogenetic trees used in the SH-test.** Nexus file can be opened in Mesquite and gene losses visualised with the “trace character” option.

### A burst of gene losses occurred in the Afrotherian stem lineage

Our observations that gene loss data likely has a strong phylogenetic signal suggest we can use evolutionary methods to reconstruct ancestral gene losses in *Atlantogentata*. Therefore, we used the Dollop program in Phylip to infer ancestral gene loss events in each branch of the species phylogeny under the Dollo model (**Figure 4A**). While there was no correlation between branch length, expressed as time since divergence in millions of years, and the number of gene loss events per branch (Pearson r^2^=0.10, *P*=0.61), several branches had more genes losses than expected, including Talazac’s shrew tenrec, Southern three-banded armadillo, and the Afrotherian stem-lineage (**Figure 4B**); these data are consistent with our observation that the southern three-banded armadillo is inferred to be its own cluster by FCM and MDS. Species-specific gene losses, such as those observed in Talazac’s shrew tenrec and Southern three-banded armadillo, arise from assembly errors. Thus, we conclude that the elevated gene loss rate in Talazac’s shrew tenrec and Southern three-banded armadillo may be an artifact but that there was a burst of gene losses in the Afrotherian stem lineage.

**Figure 4.**
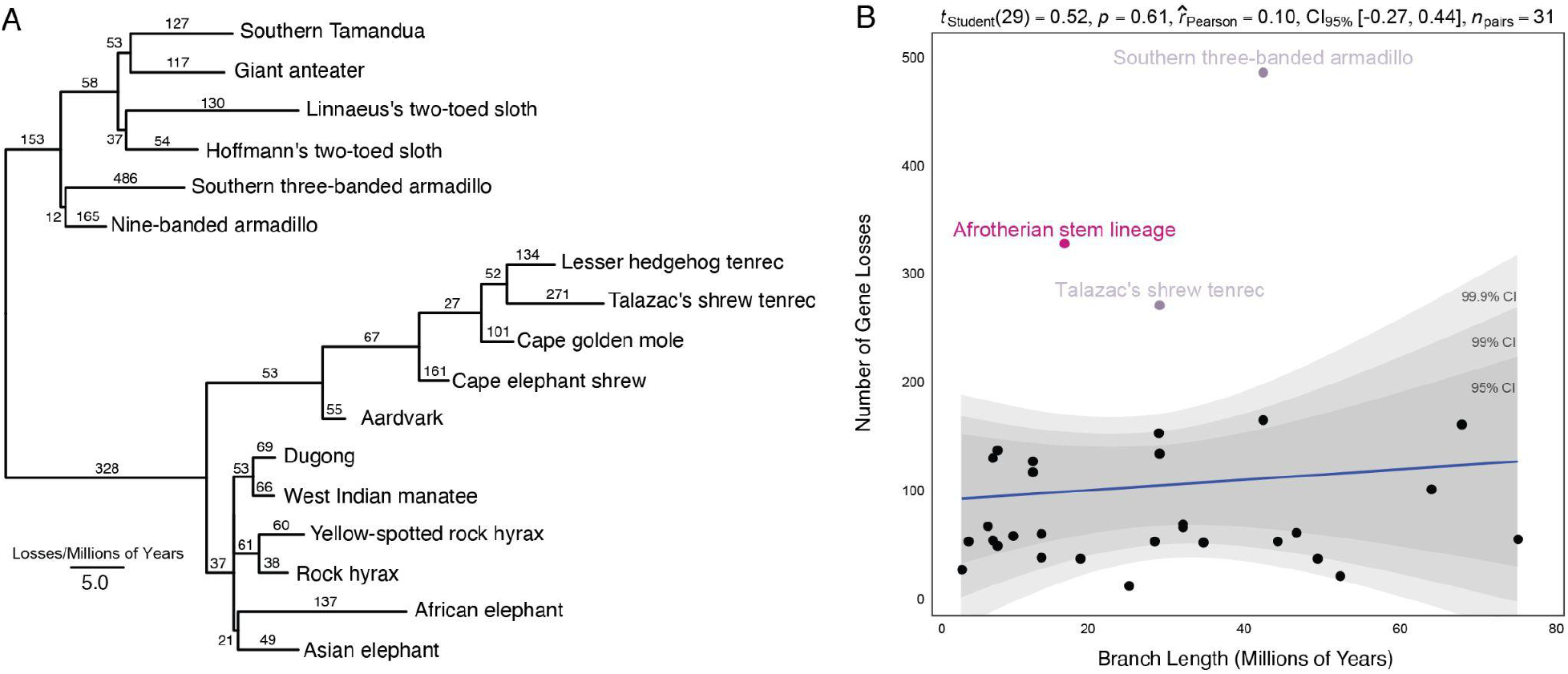
A burst of gene losses occurred in the Afrotherian stem lineage. A. Phylogenetic tree of *Atlantogenata* shows the number of gene losses inferred from the species tree and the Dollo model. Horizontal branch lengths are proportional to the rate of gene loss. The number of genes lost in each lineage is shown above each branch. Note that the relationships between elephants, hyraxes, and sea cows was collapsed to a polytomy to reflect nearly perfect uncertainty of the branching order of these lineages (Bowman 2023). B. Regression of the number of gene losses against branch length. Pearson’s correlation is shown as a blue line; 95%, 99%, and 99.9% confidence intervals (CI) are shown in grey. There is a weak (r2=0.10) but not statistically significant (P=0.61) correlation. Outlier lineages are shown in pink. **Figure 4 – source data 1.** Matrix of gene losses by branch.

### Pervasive loss of regulated cell death genes in Paenungulates

Next, we tested if lost genes were enriched in Reactome pathways (Jassal et al., 2020) and subcellular compartments (Binder et al., 2014) using the hypergeometric over-representation analyses (ORA) method implemented in WebGestalt (Liao et al., 2019). Lost genes were enriched in eight pathways and three subcellular compartments with a hypergeometric *P*≤0.05 and an FDR≤0.10. Notably, 50% (4/8) of the enriched pathways were related to regulated necrotic cell death; however, only four Reactome pathways and two cellular compartment terms remained after affinity propagation redundancy reduction, including “Regulated Necrosis” and “Inflammasome Complex” (**Figure 5A**). Lost genes were also enriched in protein-protein interaction network clusters in STRING (Szklarczyk et al., 2022) related to pyroptosis, TRAIL-activated apoptotic signaling, and the execution phase of necroptosis (**Figure 5B**). The lost genes include *MLKL* and *RIPK3*, which directly interact with each other and play critical roles in tumor necrosis factor (TNF)-induced necroptosis and have been lost in *Paenungulates*, and multiple sensor proteins that form inflammasomes and induce pyroptosis, including *AIM2* in *Afrotheria*, *MEFV* in elephants, *NLRC4* in Paenungulates and *NLRP6* in hyraxes (**Figure 5C**). In addition, *FASLG,* a transmembrane protein in the tumor necrosis factor (TNF) family expressed by immune cells, including T-cells and natural killer cells, which induces apoptosis upon binding its receptor *FAS* on target cells, has been lost in tenrecs, and several genes in the tumor necrosis factor receptor complex which inhibit cell death have been lost in many species but most conspicuously in the *Pilosa*.

**Figure 5.**
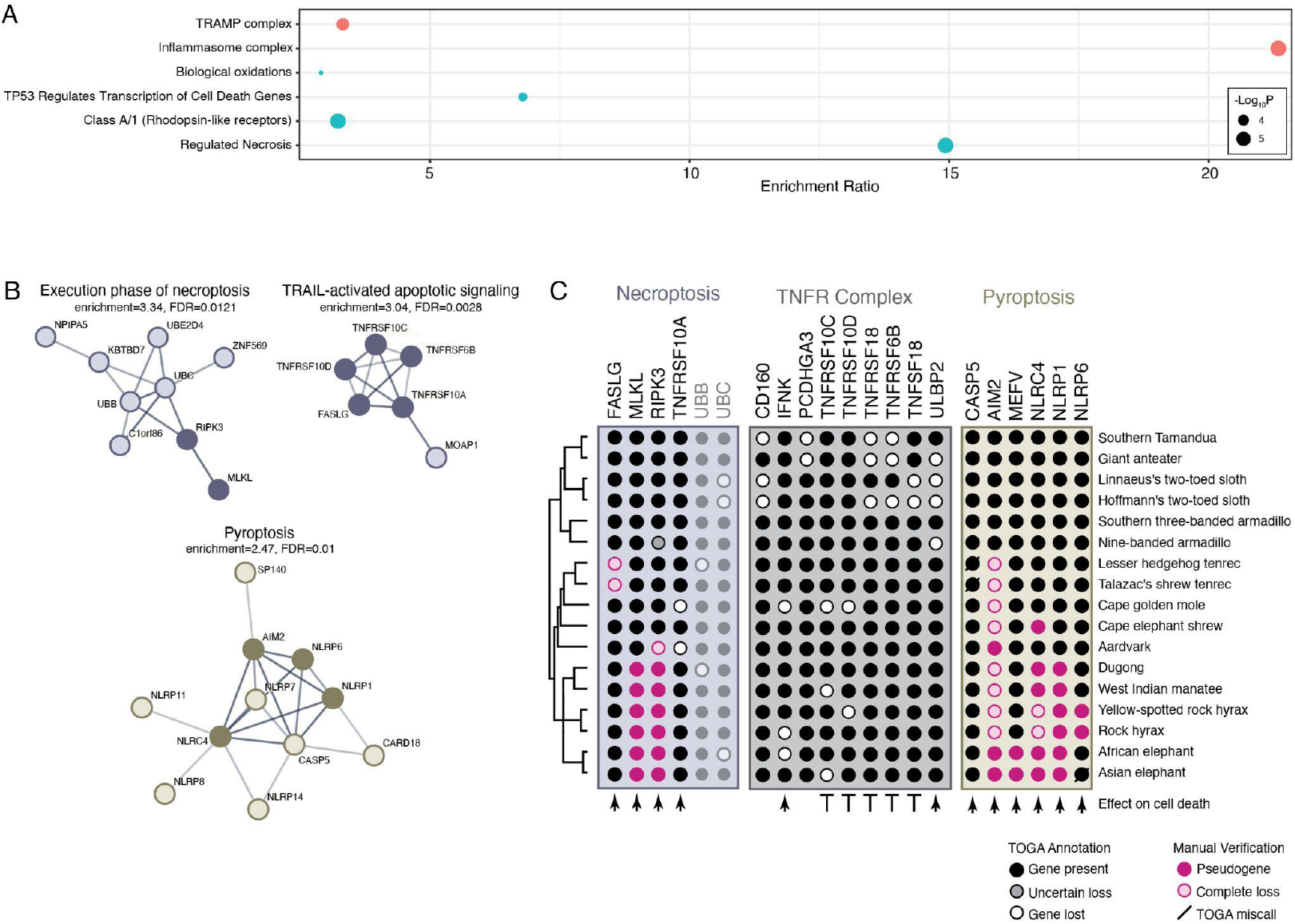
Pervasive loss of regulated cell death genes in *Afrotheria*, particularly Paenungulates. A. Lost genes are statistically enriched in subcellular compartments (Binder et al., 2014) and Reactome pathways (Jassal et al., 2020) related to regulated cell death modes; bubbleplot shows the enrichment ratio and –log_10_ hypergeometric *P*-value of enriched terms (FDR≤0.10). B. Lost genes are enriched in protein-protein interaction network clusters in STRING related to pyroptosis, TRAIL-activated apoptotic signaling, and the execution phase of necroptosis (FDR≤0.10). STRING interaction networks show protein-protein interactions with a line, and lost genes are shown with filled circles. C. Gene losses in Afritheria. Phylogenetic relationships between species are shown on the right. Genes are grouped into those that function in necroptosis (blue), the tumor necrosis factor receptor (TNFR) complex (grey), and pyroptosis (yellow). Proteins that promote cell death are indicated with arrowheads, and those that inhibit cell death are shown with blunt arrows (┴). Bubbleplot is colored by TOGA annotation and manual verification. **Figure 5 – source data 1.** Subcellular compartment enrichment results. **Figure 5 – source data 1.** Reactome pathway enrichment results.

## Discussion

Elephants have evolved remarkably large body sizes and a low prevalence of cancer (Abegglen et al., 2015; Bulls et al., 2022; Compton et al., 2023; Tollis et al., 2021). Among the anti–cancer cellular phenotypes in elephants are cells that induce apoptosis at low levels of DNA damage (Abegglen et al., 2015; Sulak et al., 2016a; Vazquez et al., 2018), are resistant to oxidative stress-induced cell death (Gomes et al., 2011), have faster DNA damage repair rates than smaller-bodied species (Francis et al., 1981; Hart and Setlow, 1974; Promislow, 1994), are resistant to experimental immortalization (Fukuda et al., 2016; Gomes et al., 2011), and that may have a genetic and/or epigenetic barrier to experimental induction of pluripotency (Appleton et al., 2024). These cellular traits are at least partly mediated by an increase in the number of tumor suppressors in the elephant lineage (Caulin et al., 2015; Doherty and Magalhães, 2016; Sulak et al., 2016b; Tollis et al., 2020; Vazquez et al., 2018; Vazquez and Lynch, 2021). Here, we explored whether the loss of oncogenes may also have contributed to cancer resistance and these anti-cancer cellular phenotypes in elephants. Unexpectedly, we found that rather than the loss of oncogenes in the elephant lineage, regulated necrotic cell death genes and pathways have been lost in many Afrotherian lineages, most notably Paenungulates. While elephants are the largest extant Paenungulate, many recently extinct Paeunungulates also evolved very large bodies, including Stellar’s sea cows, which are estimated to have weighed 10,000-11,000 kg (Scheffer, 1972), and giant hyraxes like *Titanohyrax*, which are estimated to weigh 600-1,300 kg (Tabuce, 2016). If loss of these cell death pathways protects from cancer, these data suggest that at least some anti-cancer traits evolved in the Paeunungulate stem lineage and were, therefore, permissive for the independent evolution of gigantism Paeunungulates.

### Loss of necroptosis genes in *Paenungulata*

Necroptosis is a lytic and inflammatory mode of regulated cell death regulated by RIPK3 and the MLKL (Ye et al., 2023). We found that *RIPK3* and *MLKL* have become pseudogenes in Paenungulates and that *RIPK3* is independently pseudogenized in aardvark. Our results are similar to a previous study, which found that *RIPK3* and *MLKL* have been lost in African elephants and manatees (Águeda-Pinto et al., 2021); however, Águeda-Pinto et al. only included five Afrotherians and no Xenarthrans. Thus, the extent and lineage specificity of the gene losses were unclear; our results revise and extend these observations. Remarkably, *RIPK3* and *MLKL* were also lost in other famously cancer-resistant species, including naked and blind mole rats and Cetaceans (Águeda-Pinto et al., 2021), while *MLKL* has been lost in Carnivores (Dondelinger et al., 2016; Newton and Manning, 2015). MLKL-dependent/RIPK3-independent and MLKL-independent/RIPK3-dependent necroptosis pathways have been identified (Günther et al., 2016; Zhang et al., 2016), suggesting the loss of both *MLKL* and *RIPK3* in Paenungulates and other cancer-resistant species has lead to the complete loss of this inflammatory cell death mode.

What might be the consequences of losing necroptosis? Mice with constitutively active Mlkl have normal embryonic development but develop lethal neonatal inflammation and hematopoietic dysfunction (Hildebrand et al., 2020). In contrast, *Mlkl* knockout mice cannot induce necroptosis and are protected from low-grade, sterile inflammation late in life (Crutchfield et al., 2023), chronic inflammation of the central nervous system (neuroinflammation) during normal aging (Thadathil et al., 2021), neuroinflammation and motor deficits in the α-synuclein transgenic mouse model of Parkinson’s disease (Geng et al., 2023), an age-related reduction of hematopoietic stem cells and ineffective hematopoiesis in myelodysplastic syndrome (Yamada et al., 2022), and inflammatory acute pancreatitis (Peng et al., 2023). The male reproductive tract of *Ripk3* and *Mlkl* knockout mice also retain a more ‘youthful’ morphology and function into advanced age compared to wild-type males (Li et al., 2017). These data suggest that loss of *MLKL* and/or *RIPK3* may reduce aging-associated inflammatory disease, supporting a model of aging in which inflammation contributes to the pathogenesis of age-related diseases, i.e, inflammaging (Franceschi and Campisi, 2014) as well as extend male reproductive lifespan.

Loss of necroptosis may also directly protect against cancer and metastasis. Necroptosis of tumor cells, for example, promotes metastasis (Chen et al., 2021; Yamamoto et al., 2023; Yan et al., 2022; Yang et al., 2019), and blocking necroptosis inhibits metastasis in mouse breast cancer models (Baik et al., 2021; Jiao et al., 2018; Liu et al., 2023). Similarly, inhibiting necroptosis reduced tumor-cell-induced endothelial necroptosis, tumor cell extravasation, and metastasis (Strilic et al., 2016). Loss of *MLKL* may also protect against immune evasion in hepatocellular carcinoma by suppressing parthanatos (Jiang et al., 2023), a cell death mode that occurs following the overactivation of the DNA repair enzyme poly(ADP-ribose) polymerase 1 (PARP1) (David et al., 2009; Yu et al., 2006). Unexpectedly, the deletion of *RIPK3* in mouse embryonic fibroblasts dramatically suppressed reprogramming into induced pluripotent stem cells (iPSCs), likely by reducing the expression of cell cycle progression genes (Al-Moujahed et al., 2019). Necroptosis likely functions as a backup in case of apoptotic failure (Ye et al., 2023); the loss of *RIPK3* and *MLKL* may direct cells away from inflammatory necroptotic cell death to apoptosis. Thus, loss of *MLKL* and the likely necroptosis may have multiple cancer-protective effects, while loss of *RIPK3* might act as a barrier to the induction of pluripotency.

### Loss of NET formation genes in *Paenungulata*

Neutrophils (and other granulocytes) can induce a form of inflammatory regulated cell death called NETosis, in which necrosis is accompanied by the release of neutrophil extracellular traps (NETs), characterized by intracellular degranulation and the release of decondensed chromatin and granular contents into the extracellular space (Metzler et al., 2011; Papayannopoulos et al., 2010; Vorobjeva and Chernyak, 2020). In addition to executing necroptosis, RIPK3 and MLKL mediate NET formation in necroptotic neutrophils, which release NETs at membrane locations surrounded by MLKL (D’Cruz et al., 2018); however, while knockout of *RIPK3* and *MLKL* prevents NET formation it does not prevent cell death because of residual caspase-8 activity (D’Cruz et al., 2018), which is the molecular switch that governs regulated cell death by either apoptosis, necroptosis, or pyroptosis (Fritsch et al., 2019). Remarkably, we previously found that genes related to neutrophil biology were positively selected or rapidly evolving in elephants, including genes that mediate neutrophil extracellular trap formation (Bowman and Lynch, 2024). While neutrophils have well-characterized roles in the innate immune system, they also play an important role in cancer biology, where they can both promote and inhibit tumor initiation, growth, and metastasis (Coffelt et al., 2016). These data suggest neutrophil phenotypes related to cancer biology may be derived in Paenungulates. Unfortunately, few studies have functionally characterized Paenungulate neutrophils, but some data indicate that neutrophil biology is different in elephants than in other species; elephants have higher concentrations of neutrophils in their blood than expected from their body mass (Downs et al., 2019), neutrophils that are activated and produce oxygen radicals at much lower stimuli than other species (Smith et al., 1998; Tell et al., 1999). Further studies are needed to determine if these phenotypes are unique to elephants or common for Paenungulates, if NET formation is different in Paenungulates than other species, and if Paenungulates cells are resistant to NETosis.

### Loss of multiple inflammasome sensor genes in *Afrotheria*

Pyroptosis is a highly inflammatory mode of regulated cell death that occurs in response to infection with intracellular pathogens such as bacteria, viruses, and other pathogen-associated and non-biological danger/damage-associated molecular patterns (Wei et al., 2022). There are multiple sensor proteins involved in canonical pyroptosis, each of which recognizes distinct molecules (Martinon et al., 2002; Rathinam et al., 2012; Wei et al., 2022): 1) NLRP3, which forms an inflammasome in response to endogenous danger signals, such as pore-forming toxins, crystalline structures, and extracellular ATP or RNA (Elliott and Sutterwala, 2015; Lamkanfi and Dixit, 2014); 2) AIM2, which forms an inflammasome in response to directly binding to cytosolic double-stranded DNA (dsDNA) from DNA viruses or self DNA (Fernandes-Alnemri et al., 2010; Hornung et al., 2009); 3) NLRP1, which forms an inflammasome in response multiple triggers, including anthrax LeTx toxin, muramyl dipeptide (a component of bacterial peptidoglycan), dsRNA, and enteroviral 3 C protease (Bauernfried et al., 2021; Fink et al., 2008; Rathinam et al., 2012); 4) PYRIN (encoded by *MEFV*) which forms an inflammasome in response to pathogenic toxins such as cytotoxic TcdB (Xu et al., 2014); 5) NLRC4, which forms an inflammasome in response to bacterial type III secretion system apparatus components flagellin or PrgJ (Miao et al., 2010); and 6) NLRP6, which forms an inflammasome in response to lipoteichoic acid, a major component of gram-positive bacterial cell walls (Shen et al., 2021). When these sensors are stimulated, they induce inflammasomes to produce mature caspase-1, which cleaves GSDMD, leading to pore formation, and IL-1β and IL-18 cytokine release, and an inflammatory response (Wei et al., 2022). Our observations that *AIM2* has been lost in *Afrotheria*, *NLRP1* and *NLRC4* have been lost in Paenungulates, *MEFV* has been lost in elephants, and *NLRP6* has been lost in hyraxes suggest two scenarios: 1) these lineages evolved alternative ways of activating pyroptosis in response to dsDNA, bacterial toxins and peptidoglycan, bacterial type III secretion systems, and pathogenic toxins, respectively, that rendered these sensors redundant; or 2) major canonical pyroptosis pathways have been lost in these lineages.

What may be the consequences of losing these inflammasomes? Impairment of the default CASP1/GSDMD-dependent pyroptosis pathway induced by activation of the NLRC4 inflammasome initiates an alternative ASC-mediated caspase–8–dependent apoptosis pathway while blocking both triggers intrinsic apoptosis (Zhang et al., 2021). Thus, loss of the AIM2-, NLRP1-, NLRC4-, PYRIN-, and NLRP6-inflammasomes may shift cells from pyroptosis, a highly inflammatory cell death mode, towards anti-inflammatory apoptotic cell death. Loss of *NLRP1*, for example, may contribute to reduced cancer risk by limiting inflammasome activation and prompting apoptosis, as has been observed in human metastatic melanoma (Zhai et al., 2017) and several small molecule inhibitors of the NLRP3 inflammasome are in clinical trials to block pyroptosis and suppress chronic inflammation (Coll et al., 2022). Other species have also lost these genes, including carnivores that independently pseudogenized or completely lost NLRP1 and NLRC4 (Ahn et al., 2016; Digby et al., 2021) and bats that have independently lost AIM2 (Ahn et al., 2016) and evolved dampened ASC2-mediated inflammatory responses (Ahn et al., 2023). These data suggest that the loss of some inflammasomes can be tolerated and may have adaptive benefits, such as reduced inflammation.

### Caveats and limitations

Our observation that genes important for mediating necroptosis, NET formation during NETosis, and different kinds of inflammasome-induced pyroptosis suggest that these cell death pathways have been lost in several Afrotherian lineages, most notably in Paenungulates, which have independently evolved large bodied lineages. While we have interpreted these gene loss data within the context of cancer biology and the evolution of large bodies, the loss of these genes may have had other effects or no effect on the evolution of cancer resistance or body size evolution. It is possible that species that have lost these genes have evolved alternative ways to induce necroptosis, NET formation, and inflammasome activation and that developmental systems drift has changed the roles these genes play in normal development, minimizing the effects of their loss through developmental systems drift (Lynch, 2009; True and Haag, 2001). We also cannot infer that the direction of causation, for example, was loss of these genes adaptive or was their loss a consequence of relaxed selection on these cell death pathways, relaxed selection on the functions of these genes, and their loss was a non-adaptive process? Finally, functional studies are required to determine if cells from Paenungulates are resistant to necroptotic cell death, if neutrophils have impaired NET formation, and if immune cells are resistant to the signals that induce pyroptosis in response to AIM2, pyrin, NLRC4, NLRP1, and NLRP6 activation.

## Conclusions

Our observations that genes essential for inducing different modes of regulated necrotic cell death have been lost in Afrotherians (*AIM2*), particularly the Paenungulate stem lineage (*RIPK3*, *MLKL*, *NLRC4*, *NLRP6*), suggest that these lineages have lost necroptosis, NET formation during NETosis, and some forms of pyroptosis. Remarkably, necrotic cell death modes have repeatedly been lost across the mammalian tree, including the loss of *RIPK3* and *MLKL* in naked and blind mole rats, lagomorphs, and Cetaceans (Águeda-Pinto et al., 2021), the loss *MLKL* in Carnivores (Dondelinger et al., 2016; Newton and Manning, 2015), and the loss of *NLRP1* and *NLRC4* inflammasomes in carnivores (Ahn et al., 2016; Digby et al., 2021), and *AIM2* inflammasomes in bats (Ahn et al., 2016). These data suggest that the loss of these genes may contribute to the evolution of large bodies, long lifespans, and cancer resistance in many mammalian lineages, which may have come at a cost of increased susceptibility to viral, bacterial, and other pathogens. Finally, our data indicate that there is significant turnover of regulated cell death modes in mammals, and important insights into the basic biology of cell death and immunology can be gained from studies of non-model organisms (Minton, 2023).

## Acknowledgments

This study was supported by NIH award 1R56AG071860-01 to VJL and a National Center for Case Study Teaching in Science REU to AS.

## Methods

### Phylogenetic analyses of gene loss data

We downloaded TOGA loss_summ_data.tsv files, which used the human hg38 reference for each species, from https://genome.senckenberg.de/download/TOGA/. Gene losses in each species were identified as genes (GENE) annotated as “clearly lost” or “L” and were coded as state 0, while all other TOGA categories were collapsed into a likely present category and coded as state 1; we note that this coding is conservative, as genes annotated with an “uncertain loss” and “partially intact” may be true losses. However, this coding scheme ensures we only infer losses with the highest confidence of true loss and may miss recent pseudogenization events when there are only one or a few inactivating mutations. We manually confirmed the presence/absence of functional *RIPK3*, *MLKL*, *FASLG*, *AIM2*, *MEFV*, *NLRC4*, *NLRP1*, and NLRP6 genes using BLAT search to the genome of each species in which TOGA inferred a loss.

### Exploratory data analyses

We used Multi-Dimensional Scaling (MDS) and Fuzzy C-Means (FCM) clustering to explore if species clustered by phylogenetic relationships, as expected if gene loss data had phylogenetic signal, or randomly, as expected if gene loss data was mostly noisy and TOGA had a high rate of false positive and/or losses were the result of poor genome quality. MDS was performed using the vegan R package (Oksanen et al., 2008) with four reduced dimensions. Unlike MDS, FCM allows each sample to have membership in multiple clusters; FCM membership coefficients can account for multiple sources of similarity, including noise, phylogenetic signal, and convergence of gene losses. FCM was performed in R using the fanny in the cluster package, using Manhattan distances (cluster membership was not altered by using other distance metrics), and an estimated fuzzifier (m=1.0). FCM clustering requires a priori knowledge of the number of clusters (K) to include; therefore, we evaluated FCM with K=2–10. First, we used the “elbow” method, in which the sum of squares of each cluster number is calculated and graphed and the optimal number of clusters is estimated by a change of slope from steep to shallow (the elbow). We also assessed the optimal number of clusters using the clustree R package, which assesses the optimal number of clusters by considering how samples change groupings as the number of clusters increases; clustree is useful for estimating which clusters are distinct and which are unstable but cannot determine the optimal number of clusters (K).

### Phylogenetic analyses of gene loss data

We used the dollopenny program in Phylip (version 3.695) to infer the maximum parsimony phylogeny from gene loss data under the Dollo model. Branch support was assessed with 100 bootstrap replicates (generated from seqboot in Phylip) using the dollopenny program and a parsimony variant of the Shimodaira-Hasegawa test implemented in dollop to assess the support for each branch by comparing the inferred most parsimonious tree to one in which each branch is iteratively collapsed to a polytomy. Ancestral states were inferred with the dollop program in Phylip, using the species phylogeny (with Paenungulate lineages as a polytomy) and enforcing all genes as present in the common ancestor.

### Over-Representation Analyses (ORA)

We used WebGestalt v. 2019 (Liao et al., 2019) to identify enriched Reactome pathway terms (Jassal et al., 2020) and protein subcellular localizations (Binder et al., 2014) using over-representation analysis (ORA). We excluded gene families for which orthology assignments are particularly problematic (black list), including uncharacterized open reading frames (cORFs), beta-defensins (DEFBs), members of the FAM gene family, histones, keratins (KRTs) and keratin-associated proteins (KRTAPs), nuclear pore complex interacting protein family members A and B (NPIPA and NPIPB), PRAME genes, SLCs, and SPATA, USPs, and zinc finger protein family genes (ZFPs and ZSCANs), and UDP glucuronosyltransferase family members. Statistical significance of over-representation was assessed using the set of genes lost in any internal branch combined into a single gene set as the foreground, the set of all genes in the human (hg38) reference as the background gene set (n=22,073), and a hypergeometric test. Because the number of lost genes in the foreground is large (n=548), we set the minimum number of genes for a category to 10, and the maximum to 200, and the number of categories expected from the set cover was 10; this reduces the pathways we test with ORA to those with more than 20 but less than 200 genes, which reduces the likelihood of a false positive enrichment because our foreground gene set is large and covers 2.6% of the background gene set. False discovery was controlled with the Benjamini-Hochberg false discovery rate (FDR q-value), reporting only those terms with an FDR q-value ≤ 0.10. We used STRING to identify if lost genes were enriched in protein-protein interaction networks, and sub-networks with MCL clustering with an inflation parameter of 2 and false discovery controlled with the Benjamini-Hochberg false discovery rate (FDR q-value), reporting only those terms with an FDR q-value ≤ 0.10.

