## Supplementary figures and images for "Pervasive loss of regulated necrotic cell death genes in elephants, hyraxes, and sea cows (*Paenungualta*)"

### goslim_summary_wg_result1709049343.png

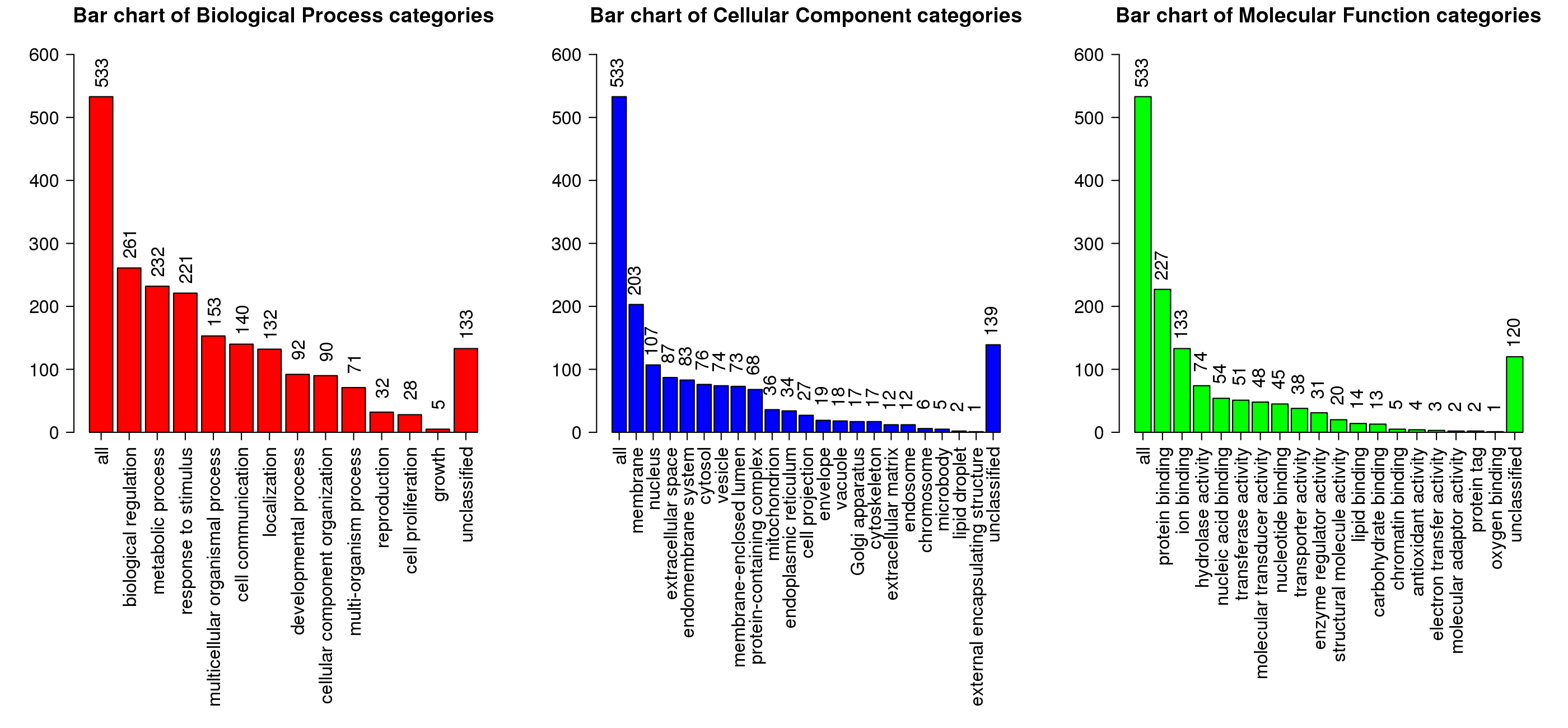
